# Ribosome dimerization prevents loss of essential ribosomal proteins during quiescence

**DOI:** 10.1101/806711

**Authors:** Heather A. Feaga, Mykhailo Kopylov, Jenny Kim Kim, Marko Jovanovic, Jonathan Dworkin

## Abstract

The formation of ribosome dimers during periods of quiescence is widespread among bacteria and some higher eukaryotes. However, the mechanistic importance of dimerization is not well understood. In bacteria ribosome dimerization is mediated by the Hibernation Promoting Factor (HPF). Here, we report that HPF from the Gram-positive bacterium *Bacillus subtilis* preserves active ribosomes by preventing the loss of essential ribosomal proteins. Ribosomes isolated from strains either lacking HPF (Δ*hpf*) or encoding a mutant allele of HPF that binds the ribosome but does not mediate dimerization were substantially depleted of the small subunit proteins S2 and S3. Strikingly, these proteins are located at the ribosome dimer interface. We used single particle cryo-EM to further characterize ribosomes isolated from a Δ*hpf* mutant strain and observed that many were missing S2, S3, or both. These data support a model in which the ribosome dimerization activity of HPF evolved to protect labile proteins that are essential for ribosome function.

**Significance Statement:** When nutrients become scarce, many bacterial species enter an extended state of quiescence. A major challenge of this state is how to attenuate protein synthesis, the most energy consuming cellular process, while preserving ribosomes for the return to favorable conditions. Here, we show that the ribosome-binding protein HPF which dimerizes ribosomes functions to protect essential ribosomal proteins at the dimer interface. HPF is almost universally conserved in bacteria and HPF deletions in diverse species exhibit decreased viability under nutrient limitation. Our data provide mechanistic insight into this phenotype and establish a role for HPF in maintaining translationally competent ribosomes during quiescence.

## Introduction

The majority of bacteria exist in a quiescent state and live in nutrient-limited conditions where conservation of resources is crucial for long-term survival (1). Protein synthesis consumes more than half of the energy in the cell during active growth (2) so when nutrients become scarce it must be tightly controlled (3). Protein synthesis is catalyzed by the ribosome, a large macromolecular complex (molecular weight of ∼2.6 million Daltons) that constitutes > 25% of the mass of the cell (4, 5). Production of new ribosomes is the most energy intensive process in the cell (6); therefore, a mechanism to preserve active ribosomes would impart a strong selective advantage during stress and nutrient limitation.

Ribosomes in quiescent cells ranging from bacteria (3, 7, 8) to some mammalian cells (9, 10) form dimers. However, the functional consequences of dimerization remain mysterious. In bacteria, ribosome dimerization is mediated by the protein HPF. Two monomers of HPF bind two 70S ribosomes and interact at their C-termini to form 100S ribosome dimers (11). Most bacteria encode an HPF homolog, and mutants lacking HPF exhibit pleiotropic phenotypes, including decreased viability during extended stationary phase (12), increased antibiotic sensitivity (13), decreased virulence (14), and a reduction in protein synthesis and growth after nutrient limitation (15, 16). Ribosome dimerization may inhibit protein synthesis (17, 18) and bacteria lacking HPF exhibit increased polysomes, indicative of increased protein synthesis (19). However, phenotypes associated with HPF deletion may also be explained by ribosome instability as ribosomal RNA has been shown to be degraded in some mutants lacking HPF (15, 20, 21).

Recent structures of 100S ribosome dimers from *Escherichia coli, Staphylococcus aureus,* and *B. subtilis* reveal that HPF binding to the ribosome would occlude binding of mRNA and initiation factors to the ribosome (11, 22, 23). In *E. coli*, the small protein RMF binds two 70S ribosomes to form 90S ribosome dimers, which are then bound by short form of HPF to produce 100S ribosome dimers (22). The short form of HPF is restricted to gamma-proteobacteria, and most bacterial species, including *B. subtilis*, encode the long form of HPF. Long HPF can form homo-dimers in solution, which are then thought to bind two 70S ribosomes to form the 100S dimer (24). Both short and long HPF bind to the same region of the ribosome and should inhibit binding of the A- and P-site tRNAs. These data therefore suggest a straightforward mechanism for inhibiting translation. However, several pieces of data suggest that HPF is not sufficient to inhibit translation *in vivo*. In particular, HPF is expressed in all growth phases in some bacteria and over-expression of HPF does not inhibit cell growth and division (25, 26).

To investigate how HPF contributes to the physiology of quiescence, we used the model Gram-positive bacterium *B. subtilis*, which contains a well-characterized HPF homolog that dimerizes ribosomes during stationary phase (11, 16). We observed that ribosomes harvested from stationary phase cells lacking HPF were severely attenuated in protein synthesis. In addition, a mutant form of HPF which binds the ribosome but does not induce dimer formation also failed to preserve the translational activity of stationary phase ribosomes. Using mass spectrometry to characterize these ribosomes, we found that two essential proteins of the small ribosomal subunit located at the dimer interface were significantly depleted in both HPF deficient mutants. Cryo-EM analysis of ribosomes harvested from stationary phase cells lacking HPF demonstrated that a substantial population of ribosomes was missing one or more essential proteins of the small subunit. Thus, our results support a model in which dimerization mediated by HPF preserves ribosomes for when conditions become favorable.

## Results

### Ribosomes harvested from stationary phase Δ*hpf* cells are inactive for protein synthesis

Ribosome dimerization mediated by HPF occurs in stationary phase in many bacteria (27), including *B. subtilis* (Fig. S1). Phenotypes of Δ*hpf* strains include a reduction in protein synthesis and a longer lag phase after nutrient limitation (15, 16). Since these phenotypes suggest that Δ*hpf* strains may have impaired translation capacity, we hypothesized that HPF is necessary to preserve ribosomes during stationary phase. To investigate this possibility, we harvested ribosomes from wild-type (JDB1772) and Δ*hpf* (JDB4221) strains in stationary phase and compared their activity by monitoring production of a reporter protein by a PURExpress *in vitro* translation system containing these ribosomes (28, 29). Ribosomes harvested from a stationary phase wild-type culture produced protein within 10 min, similar to ribosomes harvested from an exponential phase wild-type culture (Fig. 1). The stationary phase ribosomes were mainly present as dimers upon addition to the reaction and, as expected, these dimers contained HPF (Fig. S1). This experiment therefore suggests that the 100S dimer can be disassembled into free subunits that can then initiate translation *in vitro,* indicating that stoichiometric levels of HPF can be removed from the ribosome in a highly purified translation system.

**Fig. 1.**
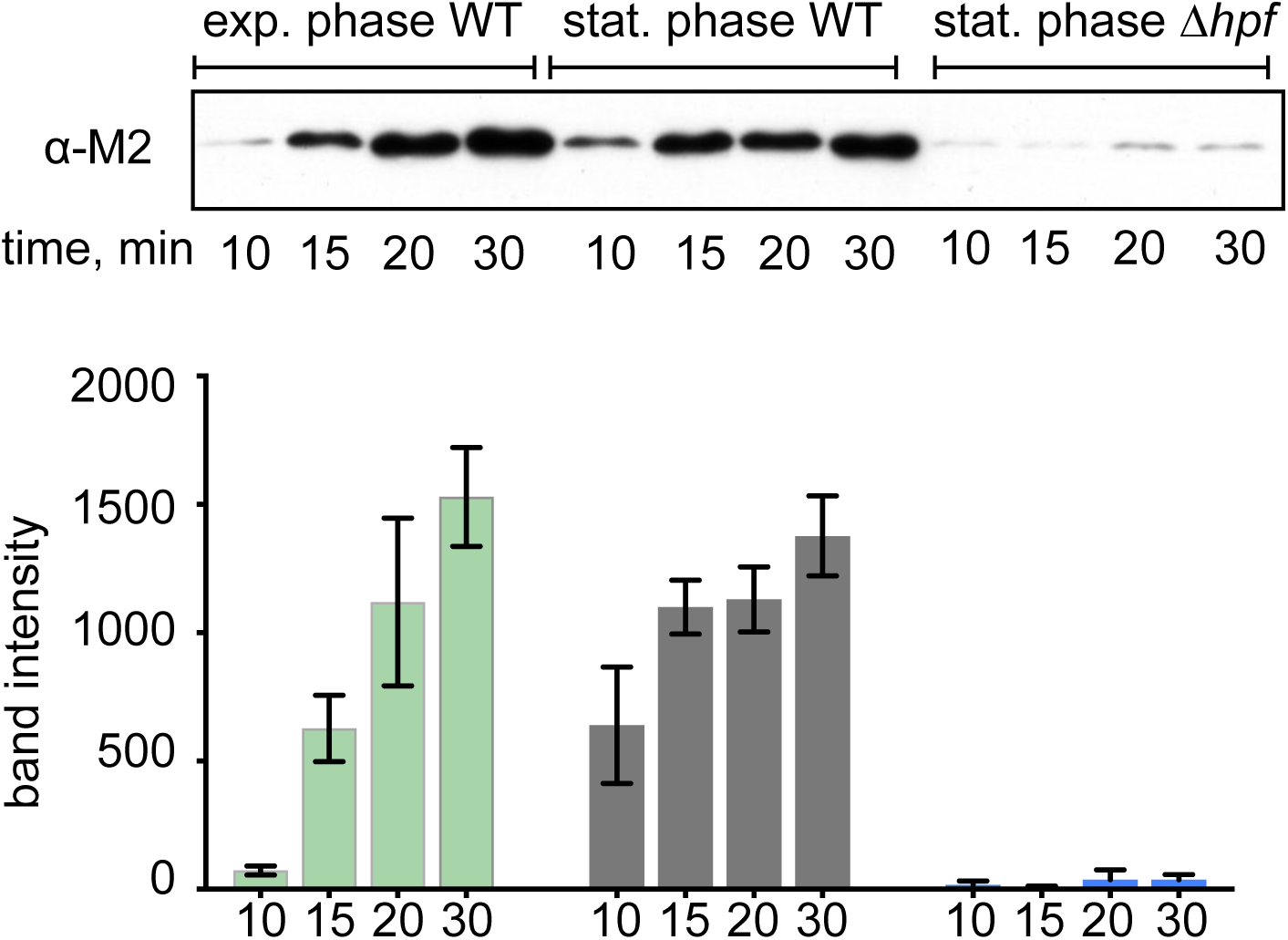
*In vitro* translation with ribosomes harvested from exponential-phase or stationary-phase cells. Ribosomes harvested from exponential phase or stationary phase wild-type cells (JDB1772), or from stationary phase Δ*hpf* (JDB4221) cells were assayed in a purified ribosome-free *in vitro* translation system. Sedimentation profiles of wild-type stationary phase ribosomes showed that these ribosomes were mainly present as dimers. Production of M2-tagged protein encoded by added template was monitored by western blot with an anti-M2 antibody (top). Reactions proceeded for 10, 15, 20, or 30 min at 37°C. Bar graph shows band intensity from 3 independent translation experiments. Error bars indicate standard deviation.

In contrast, ribosomes from a stationary-phase culture of a Δ*hpf* strain were almost entirely inactive. Protein produced by Δ*hpf* ribosomes was barely detectable even after 30 min of incubation and Δ*hpf* ribosomes exhibited 97 ± 2 % reduction in protein synthesis as compared to the wild-type stationary phase ribosomes (Fig. 1). These data suggest that HPF is required for maintaining active ribosomes.

### Ribosome dimerization is important for maintaining active ribosomes in stationary phase

The ribosome dimerization activity of HPF proteins is broadly conserved (27) but the functional consequences of dimerization are not clear. *B. subtilis* strains encoding a defective HPF that can bind the ribosome but cannot form ribosome dimers are equally defective in recovery from stationary phase as a strain lacking HPF (11). Since we observed ribosomes harvested from the HPF null strain were almost completely inactive *in vitro*, we hypothesized that the dimerization activity of HPF is required to maintain active ribosomes in stationary phase. We tested this possibility by constructing a strain expressing HPFΔdimer, an HPF mutant that binds the ribosome but is unable to form dimers. Co-migration of this mutant protein with the 70S peak indicates that it binds the ribosome but fails to mediate dimerization (Fig. 2A). We purified ribosomes from stationary phase cultures (OD_600_∼7.0) of this strain (*hpf*Δ*dimer*), as well as from wild-type and Δ*hpf* strains. Ribosomes harvested from the *hpf*Δ*dimer* strain were almost as inactive as the HPF null strain (Fig. 2B) and exhibited a 91 ± 7 % reduction in protein synthesis at 30 min compared to wild-type. Thus, ribosome dimerization, and not just ribosome binding, is necessary for preserving active ribosomes.

**Fig. 2.**
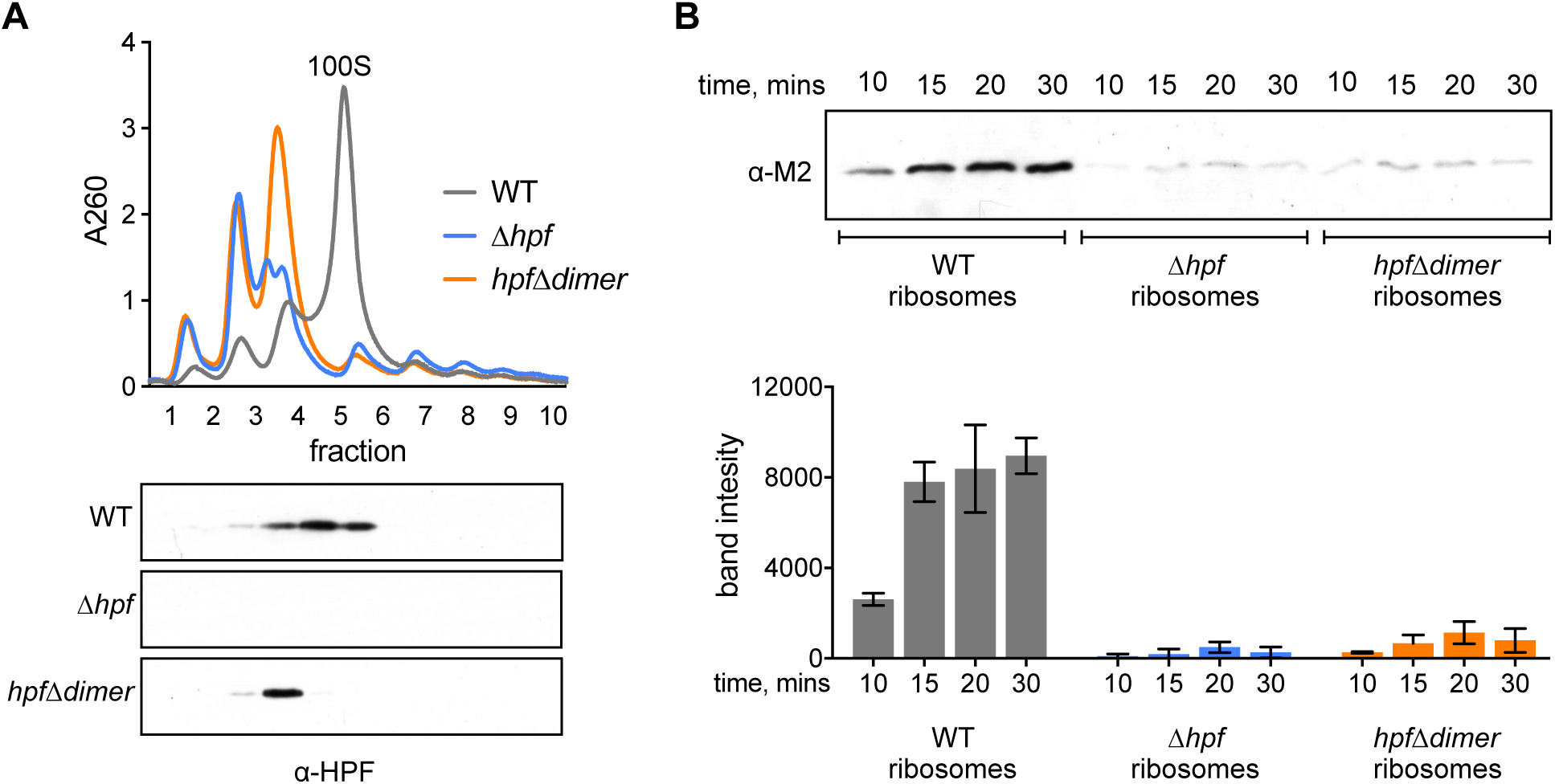
Stationary phase ribosomes harvested from HPF deficient strains are inactive. Ribosomes were harvested from wild-type (JDB1772), Δ*hpf* (JDB4221) or a strain with a mutant HPF that can bind the ribosome but does not form ribosome dimers, *hpf*Δ*dimer* (JDB4227) grown to stationary phase. (A) Sucrose density gradients of ribosomes. Fractions collected from gradients were probed with a polyclonal antibody raised against HPF. (B) Western blot with quantification from independent experiments showing translational activity of ribosomes harvested from wild-type, Δ*hpf*, or *hpf*Δ*dimer* strains. Error bars represent standard deviation.

### Stationary phase ribosomes from strains lacking functional HPF are depleted for essential proteins

While ribosomes isolated from strains lacking a functional HPF (Δ*hpf* and *hpf*Δ*dimer*) appear to maintain ribosomal RNA and most proteins (Fig. S3), they are defective in translation *in vitro* (Fig. 1, 2B) suggesting that their protein composition may be different from wild-type ribosomes. To identify potential differences, we subjected these ribosomes as well as ribosomes isolated from the wild-type cells to mass spectrometric analysis (Table 1). One protein that was more abundant in both of the mutant ribosomes is the ribosome silencing factor RsfS that binds the ribosome via ribosomal protein L14 (30) and interferes with subunit joining and modestly inhibits protein synthesis (31) (Table 1). However, deletion of *rsfS* did not rescue the inactive phenotype of Δ*hpf* ribosomes (Fig. S5) indicating that increased RsfS binding to the Δ*hpf* and *hpf*Δ*dimer* ribosomes is not sufficient to explain their inactivity.

**Table 1:**
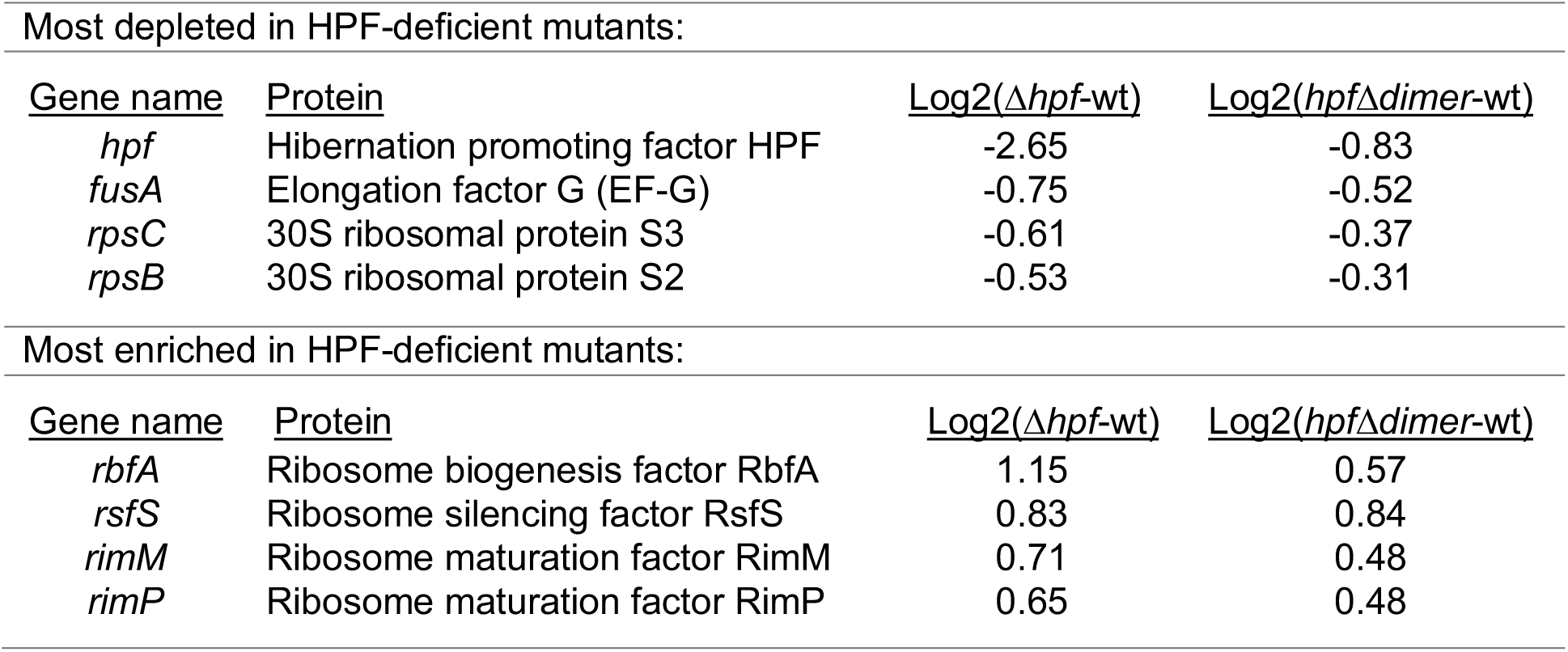
Proteins most enriched in wild-type or HPF deficient ribosomes. Values were taken from Dataset S1. The highest ranking ribosomal proteins or known ribosome interacting proteins with either positive or negative values are listed. The Log2(Δ*hpf*-wt) column is the average of two biological replicates.

A second difference between the ribosomes was that S2 and S3, two essential proteins of the small subunit, were significantly depleted in ribosomes derived from both strains relative to the wild-type (Fig. 3A; Table 1). To confirm this observation we used a polyclonal antibody raised against *B. subtilis* S2 and probed fractions collected from ribosome gradients of wild-type and the two HPF-deficient mutant strains. In ribosomes isolated from both mutant strains, S2 levels were significantly reduced, confirming the results of the mass spectrometry (Fig. 3B). Thus, the loss of S2 and/or S3 may be sufficient to explain the defect of ribosomes isolated from cells lacking a functional HPF. Strikingly, a recent structure reveals that S2 makes extensive contacts with HPF and is protected from the solvent interface when *B. subtilis* ribosomes dimerize (11).

**Fig. 3.**
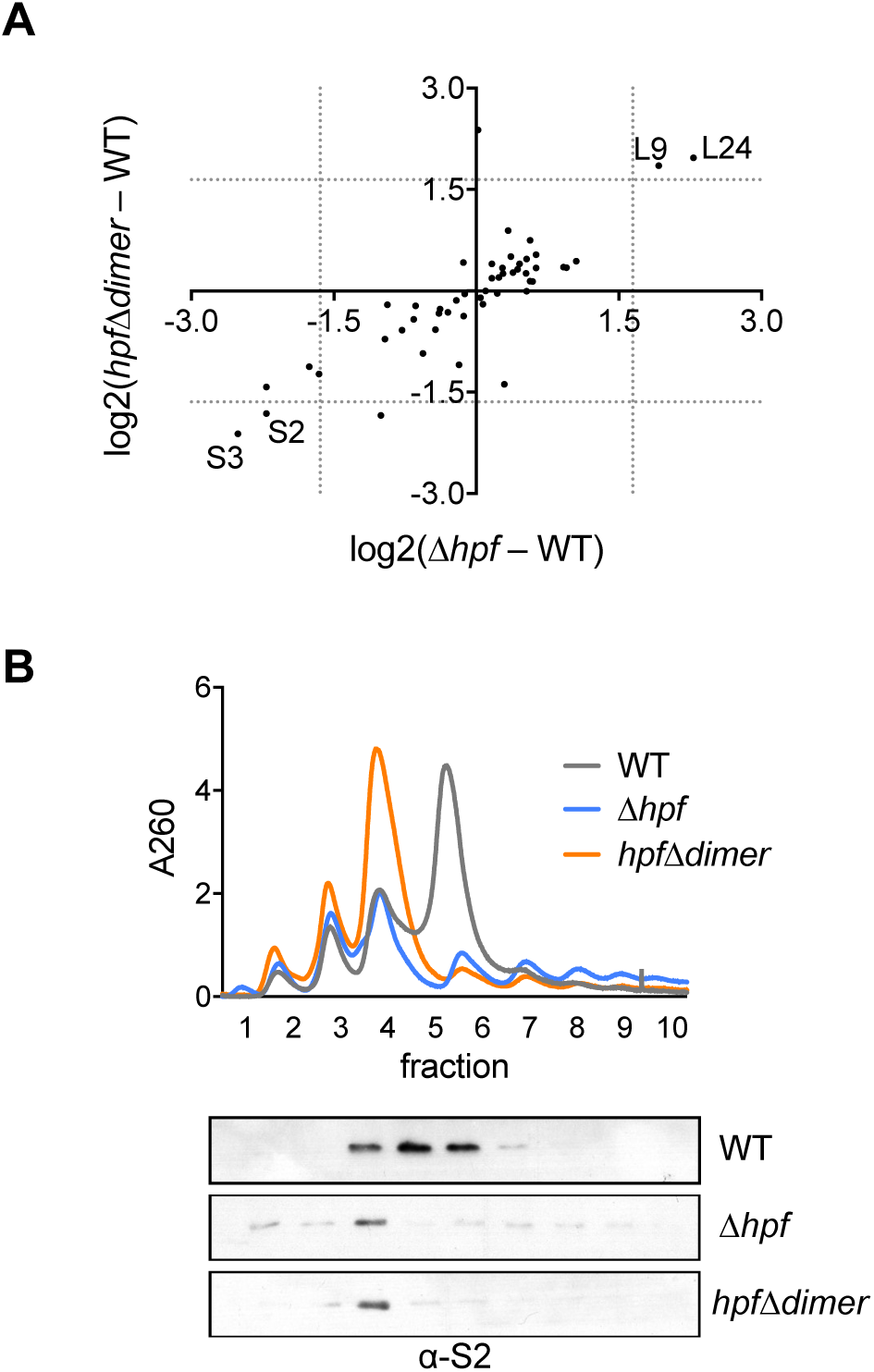
Ribosomes from strains lacking functional HPF are depleted for S2 and S3. Ribosomes were harvested from wild-type (JDB1772) cells or Δ*hpf* (JDB4221) or *hpf*Δ*dimer* (JDB4227) strains in stationary phase and analyzed by mass spectrometry and by western blot. (A) Plot of the difference in levels of ribosomal proteins between wild-type and each HPF mutant (top). Proteins in lower left quadrant are depleted in both mutants as compared with wild-type. Proteins in upper right quadrant are enriched in both mutants as compared to wild-type. Dotted lines indicate a Z-score value of 1.645. (B) Sucrose density gradients of ribosomes purified from each strain (top). Gradient fractions were probed with polyclonal antibody raised against purified S2 protein (bottom).

### Cryo-EM reveals classes of ribosomes lacking S2 and S3

Since both S2 and S3 are essential for translation, loss of only one of these proteins would be sufficient to inactivate the ribosome. We hypothesized that ribosomal subpopulations lacking only one of these proteins, or both, exist. To test this, we used single particle cryo-EM to further analyze the composition of different classes of ribosomes in the population. The Δ*hpf* strain was grown to stationary phase and the 70S ribosomal peak was isolated from a sucrose gradient, concentrated, and analyzed by cryo-EM. We observed three abundant classes of 70S ribosomes after classification and refinement. To determine which ribosomal proteins were absent, we rigid-body fitted the model of a small subunit of the *E. coli* 70S ribosome (PDB4V4A (32)) into cryoEM maps for each of these classes. The first class, comprising 59% of the particles had prominent density in the regions of both S2 and S3. A second class of particles (25%) lacked density in the region of S2 and the third class (16%) lacked density in the region of both S2 and S3 even at higher thresholds.

## Discussion

We find that *B. subtilis* HPF contributes to ribosome preservation by protecting essential proteins found on the small ribosomal subunit at the dimer interface. In strains that lack HPF or express an HPF mutant unable to mediate ribosome dimerization, ribosomal proteins S2 and S3 were significantly depleted from stationary phase ribosomes (Fig. 3). Cryo-electron microscopy further confirmed that a significant population of ribosomes harvested from a Δ*hpf* strain in stationary phase were missing S2, S3, or both (Fig. 4). Thus, dimerization may have evolved as a mechanism to protect essential ribosomal proteins that are susceptible to loss and/or degradation (Fig. 5).

**Fig. 4.**
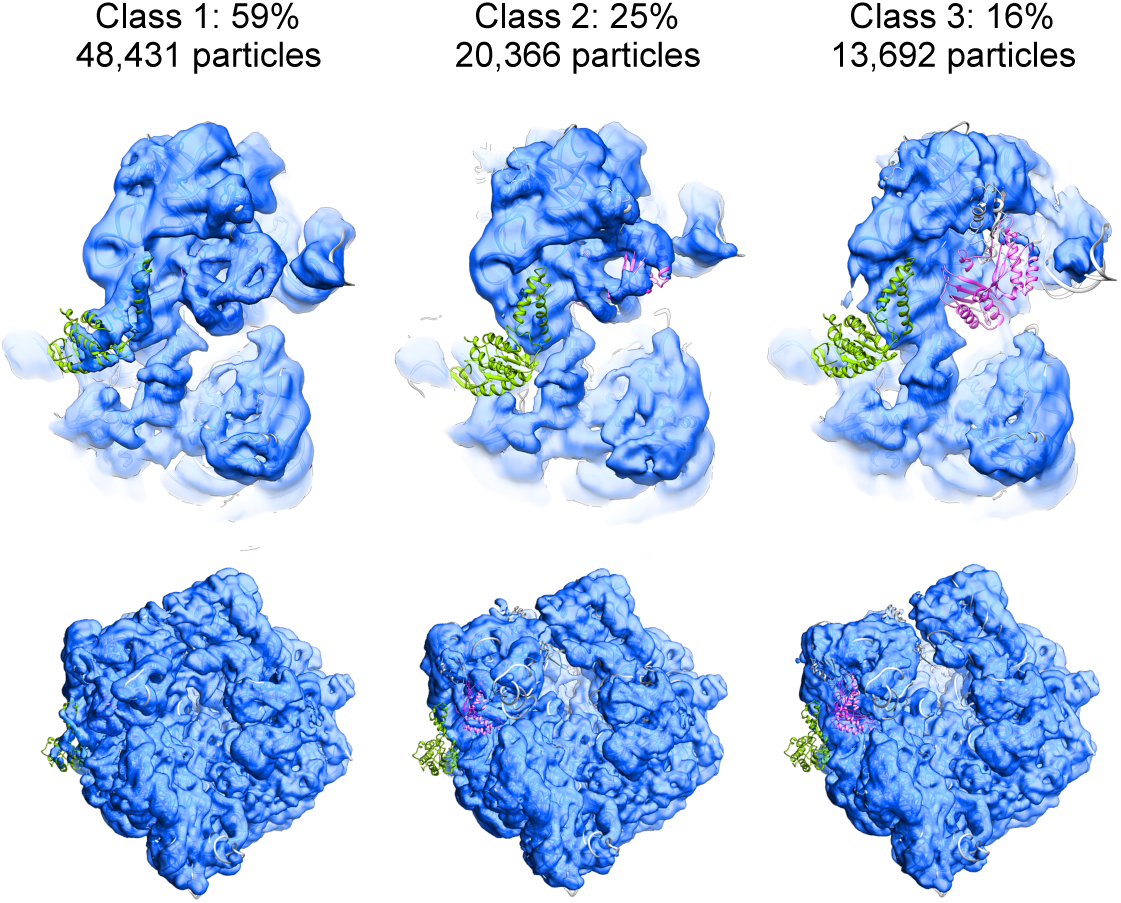
Cryo-EM analysis of Δ*hpf* ribosomes. Ribosomes were harvested from Δ*hpf* stationary phase cultures and the 70S peak was isolated by sucrose gradient centrifugation and analyzed by cryo-EM. The small subunit model from the structure of the *E. coli* 70S ribosome (ribbon diagram) was rigid-body fitted into the maps of three different classes of the Δ*hpf* ribosome (blue). Proteins S2 (green) and S3 (purple) of the *E. coli* 70S structure are indicated. Two views are shown: a zoomed-in view of the small subunit (top) and the complete 70S particle (bottom).

**Fig. 5.**
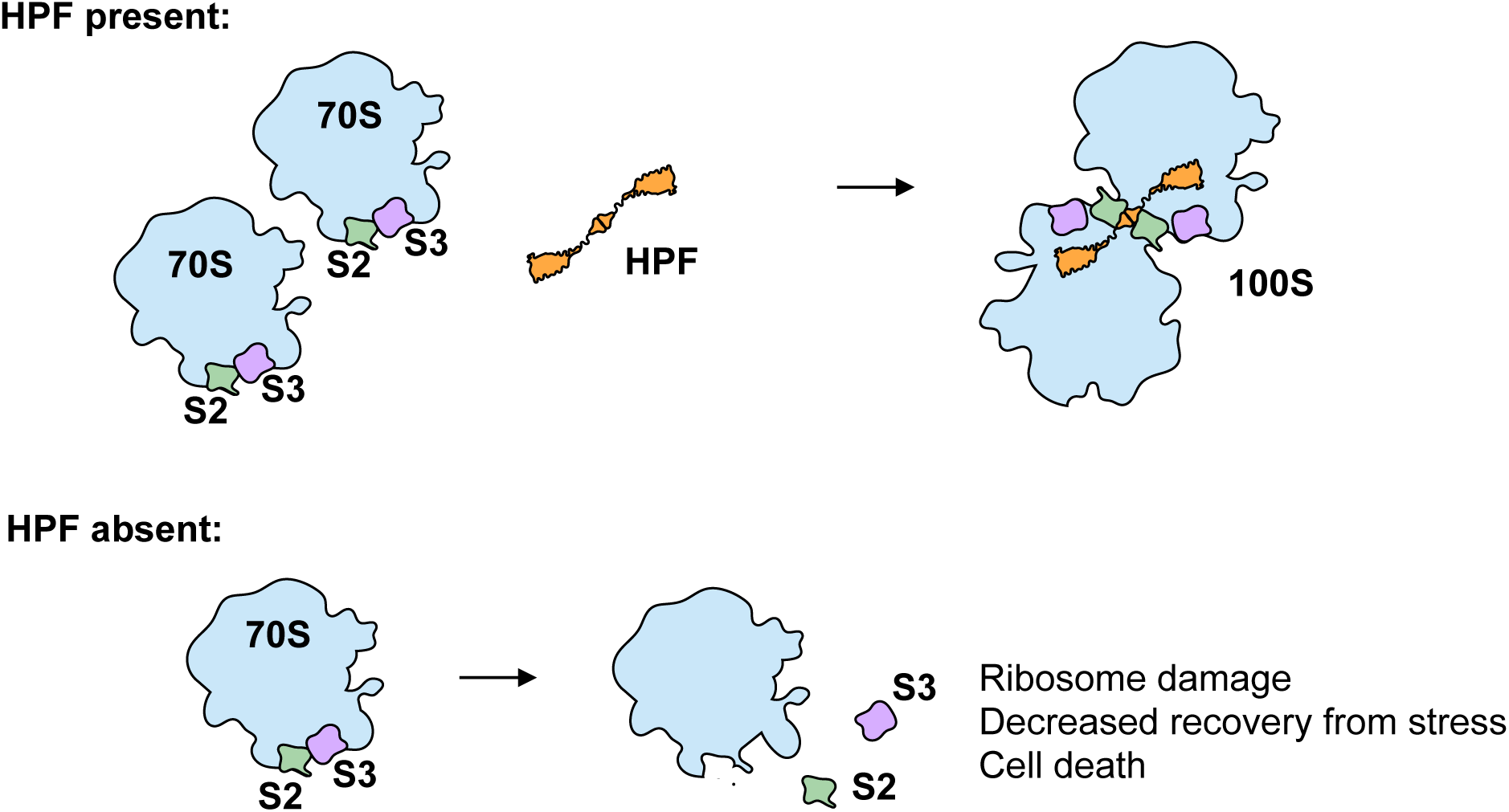
Model for HPF mediated protection of ribosomes in *B. subtilis*. HPF binds two 70S ribosomes to form a 100S dimer. S2 and S3 proteins at the dimer interface (green and purple) are protected and occluded from the solvent surface. If HPF is not functional, S2 and S3 are exposed and may not remain bound to the ribosome.

Ribosomes isolated from stationary phase cells lacking HPF are not active *in vitro* (Fig. 1). This finding may be sufficient to explain why *B. subtilis* strains lacking HPF exhibit decreased recovery from stationary phase. Interestingly, a *B. subtilis* strain expressing a mutant form of HPF that can bind the ribosome but cannot mediate dimer formation also exhibits this phenotype (16). We tested a strain expressing a similar mutant protein (*hpf*Δ*dimer*) and found that ribosomes harvested from this strain were nearly as inactive as ribosomes from the HPF null strain. We hypothesized that ribosomes harvested from either the Δ*hpf* or the *hpf*Δ*dimer* strain might be defective in similar ways. Therefore, we characterized ribosomes from both mutants by mass spectrometry (Table 1), and found that they deviated significantly from wild-type ribosomes in their protein composition (Fig. 3A, B). In particular, both mutants were significantly depleted for S2 and S3, essential proteins of the small ribosomal subunit that are located directly at the interface between the dimers. The protection of S2 is consistent with the presence of numerous direct contacts between HPF and S2; thus, in free 70S or 30S complexes, S2 is much more exposed than in the 100S dimer configuration. S3 is near the dimer interface, and participates in dimer formation in *E. coli*, but not in *B. subtilis* (11, 18). However, ribosome dimers are flexible, and as there is some movement of the 70S monomers relative to one another, S3 may also be protected by dimer formation (11, 23).

Why are S2 and S3 specifically in need of protection during stationary phase? S2 and S3 are the last proteins to be added to the ribosome during assembly and thus may easily dissociate from the ribosome (33, 34). Consistently, as much as 17% of S2 has been detected free from the ribosome in the cytosol (35). Interestingly, under stress conditions such as a shift to nutrient limitation, *E. coli* S2 is rapidly degraded by Lon protease (36). Loss of S2/S3 may further destabilize the ribosome because ribosomes not participating in translation are especially susceptible to degradation due to exposure of the rRNA when the 30S and 50S subunits are separated (37). Cryo-EM analysis resolved specific populations of ribosomes with defects in the small subunit. Approximately 40% of the ribosomes were missing either S2 or S3. Since both of these proteins are essential for translation, these data indicate that a substantial portion of the ribosomes in the Δ*hpf* strain would be inactive.

HPF is essential for preserving 16S or 23S ribosomal RNA in *E. coli, Pseudomonas aeruginosa*, and *Mycobacterium smegmatis* (15, 20, 21) but significant degradation of ribosomal RNA has not been reported in *B. subtilis* and was not observed in this study. Could loss of S2 and S3 be the first step in the degradation of ribosomes in these species? Loss of S2 and S3 and the subsequent failure to translate and form active 70S ribosomes could contribute to further destabilization of the ribosome and eventual loss of rRNA. Thus, the defective small subunit reported here may be a degradation intermediate on the way to further loss of rRNA. Additionally, failure to protect the active site of the ribosome may allow other proteins to bind which would normally be occluded such as ribosome silencing factor (RsfS) (Table 1).

Mass spectrometry analysis of stationary phase ribosomes revealed that EF-G was also deficient in the Δ*hpf* ribosomes as compared to wild-type ribosomes (Table 1). EF-G associates with the ribosome during its catalytic cycle and therefore would be expected to purify with actively translating ribosomes. Thus, its absence may further indicate that the Δ*hpf* ribosomes are inactive *in vivo*. Proteins enriched in ribosomes obtained from Δ*hpf* and *hpf*Δ*dimer* strains include the ribosome maturation factors RimM and RimP as well as RbfA that binds the 30S subunit in a position that overlaps with the A and P site tRNA binding sites similar to HPF (38). RbfA prevents initiation by a 30S subunit that has not fully matured (39, 40). Since S2 and S3 are late-binding proteins, the association of RbfA with ribosomes isolated from Δ*hpf* further indicates that the 30S may be structurally similar to a small subunit that has not completed assembly and which should be prevented from initiation. Interestingly, mutations in *E. coli* RimM lead to the loss of both S2 and S3 from immature 30S subunits (41). Similarly, ribosomes isolated from *E. coli* strains lacking RimP are depleted for S2, S3 and S21 (42). Whether the binding of these proteins is a consequence of S2 and/or S3 depletion will be a subject for future studies.

Given that ribosome synthesis is the single most energy consuming process in cells, preventing ribosome degradation, even when protein synthesis levels are low, would be evolutionarily favorable (6). In addition, a readily available, excess store of ribosomes under nutrient limitation would facilitate rapid resumption of growth when nutrients become available (43). The ability to transition to maximal protein synthesis as quickly as possible imparts a clear selective advantage for cells with higher ribosomal content (44, 45). Consistent with this interpretation, stationary phase *B. subtilis* cells lacking HPF (16) or expressing a mutant HPF unable to dimerize ribosomes exhibit both decreased viability during stationary phase and an increased lag phase following transfer to fresh media (11, 16). The long-term protection of excess ribosomes depends specifically on the ability of the cell to store these ribosomes in the form of dimers. Data presented here suggest this is because proteins located at the interface between the dimers are especially susceptible to being lost from the ribosome. Strikingly, even bacteria that encode a shortened form of HPF unable to mediate dimerization also express an accessory protein (e.g., *E.coli* RMF) that mediates dimerization. Taken together, these results suggest that ribosome dimerization evolved to guard against the loss of essential proteins of the small subunit.

## Materials and Methods

### Strains and media

Strains were derived from *B. subtilis* 168 *trpC2* and grown in LB. For detailed information on strain construction see supplemental material.

### Sucrose gradient density centrifugation

Cells were lysed using a Fast Prep machine in ribosome gradient buffer (20 mM Tris-acetate [pH_4°C_ 7.4], 60 mM ammonium chloride, 7.5 mM magnesium acetate, 6 mM β-mercaptoethanol, 0.5 mM EDTA). Lysate was cleared of debris by centrifugation at 4°C at 20,000 rcf for 20 min. Ribosomes were purified by sucrose cushion (37.7% sucrose in ribosome gradient buffer). A260 for lysates or purified ribosomes was determined by Nanodrop, and 100 μL of normalized lysate or ribosomes (as indicated in figure legends) was loaded on a 10% – 40% sucrose gradient in ribosome gradient buffer. Gradients were run for 3 hours at 30,000 rpm in a Beckman SW41 rotor. Samples were collected with a BioComp gradient station and a BioComp TRiAX flow cell monitoring continuous absorbance at 260 nm.

### *In vitro* translation

Ribosomes for *in vitro* translation were harvested by loading cleared cell lysate on a 1.5 mL buffered sucrose cushion (20 mM Tris-acetate [pH_4°C_ 7.4], 100 mM ammonium chloride, 10 mM Magnesium acetate, 0.5 mM EDTA, 6 mM β-mercaptoethanol, 37.7% sucrose). Ribosomes were pelleted in a Beckman TLA100.3 rotor at 85,000 rpm for 2 hours. Pellets were washed and resuspended in ribosome gradient buffer by gentle agitation at 4°C. Ribosomes were added to 100 nM to the PURExpress ribosome-free kit (New England Biolabs) (29). A template encoding an M2 tagged peptide was added to the reaction which was then resolved by SDS-PAGE and probed with an anti-M2 antibody (Sigma). Further details are described in supplemental material.

### Mass spectrometry

Protein sample processing and protein quantification by mass spectrometry was performed as described previously with minor modifications (46, 47). Raw data were analyzed with MaxQuant software version 1.6.0.16 (48) using a UniProt *Bacillus subtilis* 168 database (proteome UP000001570; downloaded March 2019).

### Cryo-Electron microscopy sample preparation and data acquisition

Ribosomes harvested from the Δhpf strain were resolved on a 10%–40% sucrose gradient as described. The 70S peak was isolated, exchanged into ribosome buffer without sucrose and concentrated with a Corning 100K cutoff PES filter to 200 nM. Sample was applied to plasma treated holey carbon grids, and vitrified in liquid ethane using an FEI Vitrobot. Images were automatically acquired using Leginon/Appion (49, 50) on a Titan Krios microscope (ThermoFisher Scientific) operated at 300 keV equipped with a K2 direct electron detector (Gatan) and a post-column energy filter (operated at 20 eV slit width). Movies were recorded in counting mode for 8s (20 ms/frame) with a total dose on a specimen level of 56.7 e^−^/Å^2^. Defocus range was set to vary between 0.7 and 2.0 μm. Raw frames were aligned and dose-weighted using MotionCor2 (51). CTF estimation was done with CTFFIND4 (52).

### Cryo-Electron microscopy data processing and heterogeneity analysis

Particle picking was done using template-based picking (FindEM) in Appion (53) using templates generated from manually picked particles. 2D classification, *ab-initio* reconstruction and hetero refinement steps as well as final refinements were done in cryoSPARC2 (54) using default settings and three classes (Fig. S4). Consensus refinement and masked classification were done with RELION3 (55, 56). Final maps are submitted to EMDB with the following accession codes:

## Acknowledgements

We thank the members of our lab for helpful discussions and comments on the manuscript. We thank Sandro Pereira for constructing the Δ*hpf* (JDB4221) strain. We thank Luke Berkowitz for generously providing access to his ribosome gradient analyzer. HF was supported by NIH F32GM122266; MJ was supported by R35GM128802; JD was supported by R01 GM114213, R21AI135427, and is a Burroughs-Welcome Investigator in the Pathogenesis of Infection Disease. Some of this work was performed at the Simons Electron Microscopy Center and National Resource for Automated Molecular Microscopy located at the New York Structural Biology Center, supported by grants from the Simons Foundation (SF349247), NYSTAR, and the NIH National Institute of General Medical Sciences (GM103310).

