## Supplemental Information for "Ribosome dimerization prevents loss of essential ribosomal proteins during quiescence"

**Supplementary Information Text**

**Materials and Methods.**

**Strain Construction**

The *hpf* null strain was constructed by amplifying a region of ~1kb upstream and downstream of *hpf* with primers SFP169 and SFP170 (upstream) and SFP181 and SFP182 (downstream). A tetracycline resistance marker was amplified with primers SFP1 and SFP2. The upstream and downstream flanking PCR products contained regions of homology to the primers used to amplify the $tet^R$ cassette. The complete construct, with *hpf* replaced by $tet^R$ was amplified using SFP169 and SFP182 using PCR overlap extension and transformed into PY79. Genomic DNA from the resulting strain was transformed into the *trpC2* background (JDB1772) and selected on tetracycline (10 µg/mL) to construct strain JDB4221. JDB4227 was constructed by amplifying *hpf* with primers HF40 and HF41 and ligating into pKL147 (2) digested with the same enzymes. An M2 tag was inserted between the NheI and SphI sites. The resulting plasmid was inserted at the *hpf* locus by Campbell integration. Strain JDB4292 was constructed by transforming JDB1772 with gDNA from the Bacillus Genetic Stock Center strain BKK25620 and selecting kanamycin. JDB4293 was made by transforming JDB4221 with gDNA from BKK25620.

**Purification of HPF for polyclonal antibody**

*hpf* was amplified from *B. subtilis* genomic DNA from strain JDB1772 with primers HF38 and HF39. The PCR product was digested with NcoI and XhoI and ligated into pET28b digested with the same enzymes. The resulting plasmid was transformed into BL21(DE3) to construct JDE3069 and protein expression was induced with 1 mM IPTG for 2 hours at 37˚C. Cells were lysed in Buffer A (10 mM Tris [pH 7.5], 30 mM NH4Cl, 2 mM MgCl2, and 5.7 mM β-mercaptoethanol.) Clarified lysate was incubated with Nickel-NTA resin for 1 hour at 4˚C and passed through a column. Resin was washed with 100 bed volumes of Buffer A with 20 mM imidazole. Protein was eluted with 300 mM imidazole in Buffer A and then dialyzed against Buffer A. Protein was stored at -80˚C with 20% glycerol. Antibody was obtained from Pocono Rabbit Farms and Laboratory using the Mighty Quick PHS approved protocol.

***In vitro* translation**

Ribosome concentrations were quantified by dissociating ribosomal subunits in low magnesium buffer and reading absorbance at 260 nM. To further verify that ribosomes were added equal concentration, ribosomal RNA was viewed by agarose gel electrophoresis 125 ng of a M2-tagged template was added for each translation reaction, and reactions were incubated at 37˚C. Reactions were stopped by adding SDS-PAGE loading buffer and heating to 95˚C for 5 minutes. Reactions were resolved on a 12% SDS-PAGE gel and protein production was determined by western blot with HRP-conjugated anti-M2 antibody (1:20,000; Sigma). Band intensity was determined with ImageJ.

**Mass spectrometry based protein quantification of the ribosome enriched sample**

Proteins were precipitated by a standard methanol-chloroform protocol. Briefly, first, 4 times the sample volume of methanol was added and the sample was vortexed. Second, 1 time the sample volume of chloroform was added and the sample was vortexed. Third, 3 times the sample volume water was added and the sample was vortexed. The sample was then centrifuged for 5 min at 13,000 g at room temperature. The upper aqueous layer was removed without perturbing the interface layer containing the protein precipitate. 4 times the sample volume of methanol was added and the sample was vortexed. The sample was centrifuged for 5 min at 13,000 g at room temperature, which pelleted the protein precipitate. The supernatant was removed carefully without perturbing the pellet and the remaining methanol was evaporated by air drying. The protein pellet was reconstituted in 15 µl urea buffer (8 M Urea, 75 mM NaCl, 50 mM Tris/HCl pH 8.0, 1 mM EDTA) and protein concentrations were determined by BCA assay (Pierce). 10 µg of total protein per sample were processed further. Disulfide bonds were reduced with 5 mM dithiothreitol and cysteines were subsequently alkylated with 10 mM iodoacetamide. Samples were diluted 1:6 with 50 mM Tris/HCl (pH 8.0) and sequencing grade modified trypsin (Promega) was added in an enzyme-to-substrate ratio of 1:50. After 16 h of digestion, samples were acidified with 1% formic acid (final concentration). Tryptic peptides were desalted on C18 StageTips according to (3) and evaporated to dryness in a vacuum concentrator. Desalted peptides were labeled with the TMT-11plex mass tag labeling reagent according to the manufacturer's instructions (Thermo Scientific) with small modifications. Briefly, 0.2 units of TMT-11plex reagent was used per 10 µg of sample. Peptides were dissolved in 30 µl of 50 mM Hepes pH 8.5 solution and the TMT-11plex reagent was added in 12.3 µl of MeCN. After 1 h incubation the reaction was stopped with 2.5 µl 5% Hydroxylamine for 15 min at 25°C. Differentially labeled peptides were mixed for each replicate (see mixing scheme below) and subsequently desalted on C18 StageTips (3) , evaporated to dryness in a vacuum concentrator and reconstituted in 20 µl of 3% acetonitrile and 0.1% formic acid.

| **Sample** | **TMT label** |
|---|---|
| wt (JDB1772) stationary phase (8.6.18 prep) | 126 |
| ∆hpf (JDB4221) stationary phase (8.6.18 prep) | 127N |
| wt stationary phase (12.31.18 prep) | 127C |
| ∆hpf stationary phase (12.31.18 prep) | 128N |
| hpf-M2 (JDB4227) stationary phase (12.31.18 prep) | 128C |
| wt log phase (8.6.18 prep) | 131C |
| ∆hpf log phase (8.6.18 prep) | 131 |

The samples were afterwards analyzed by LC-MS/MS on a Q-Exactive HF was performed as previously described with minor modifications (4). ~ 1 µg of total peptides were analyzed on an ACQUITY UPLC M-class system (Waters) coupled via a 50 cm Thermo Easy-Spray LC column (inner diameter of 75 µm, packed with 2 µm C18 bead stationary phase, 50 cm length, Thermo Fisher Scientific product # ES803A) to a benchtop Orbitrap Q Exactive HF mass spectrometer (Thermo Fisher Scientific). Peptides were separated at a flow rate of 250 nL/min with a linear 106 min gradient from 2% to 25% solvent B (100% acetonitrile, 0.1% formic acid), followed by a linear 5 min gradient from 25 to 85% solvent B. Each sample was run for 170 min, including sample loading and column equilibration times. Data was acquired in data dependent mode using Xcalibur 2.8 software. MS1 Spectra were measured with a resolution of 60,000, an AGC target of 3e6 and a mass range from 375 to 2000 m/z. Up to 15 MS2 spectra per duty cycle were triggered at a resolution of 60,000, an AGC target of 2e5, an isolation window of 1.6 m/z and a normalized collision energy of 36.

All raw data were analyzed with MaxQuant software version 1.6.0.16 (5) using a UniProt *Bacillus subtilis* 168 database (proteome UP000001570; downloaded March 2019), and MS/MS searches were performed with the following parameters: TMT11plex labeling on the MS2 level, oxidation of methionine and protein N-terminal acetylation as variable modifications; carbamidomethylation as fixed modification; Trypsin/P as the digestion enzyme; precursor ion mass tolerances of 20 p.p.m. for the first search (used for nonlinear mass re-calibration) and 4.5 p.p.m. for the main search, and a fragment ion mass tolerance of 20 p.p.m. For identification, we applied a maximum FDR of 1% separately on protein and peptide level. We required 1 or more unique peptides for protein identification and a ratio count for each of the 11 TMT channels of the corresponding TMT-11plex mix. This gave us a total of 763 quantified protein groups with at least 1 identified peptide, 450 proteins with 3 or more identified peptides and a total of 51 ribosomal proteins.

Finally, each protein group of a TMT labeled sample got its proportional fraction of the MS1 based iBAQ intensities based on its labeling channel specific TMT MS2 intensity relative to the sum of TMT MS2 intensities of all labeled channels for the corresponding protein group. Afterwards we normalized these fractional MS1 iBAQ intensities such that at each condition/time point these intensity values added up to exactly 1,000,000, therefore each protein group value can be regarded as a normalized microshare (we did this separately for each TMT channel for all proteins that made our filter cutoff in all the TMT channels of the corresponding TMT-11plex mix). After that we added a pseudocount of 1 to each intensity value in order to account for the noise level and make our fold change calls more robust for small intensity values. Finally, we log2 transformed all values and subtracted from the log2 protein values of each hpf mutant sample its corresponding wildtype sample log2 protein values, which provided the log2 ratios changes for each protein group in an hpf mutant relative to its corresponding wildtype sample.

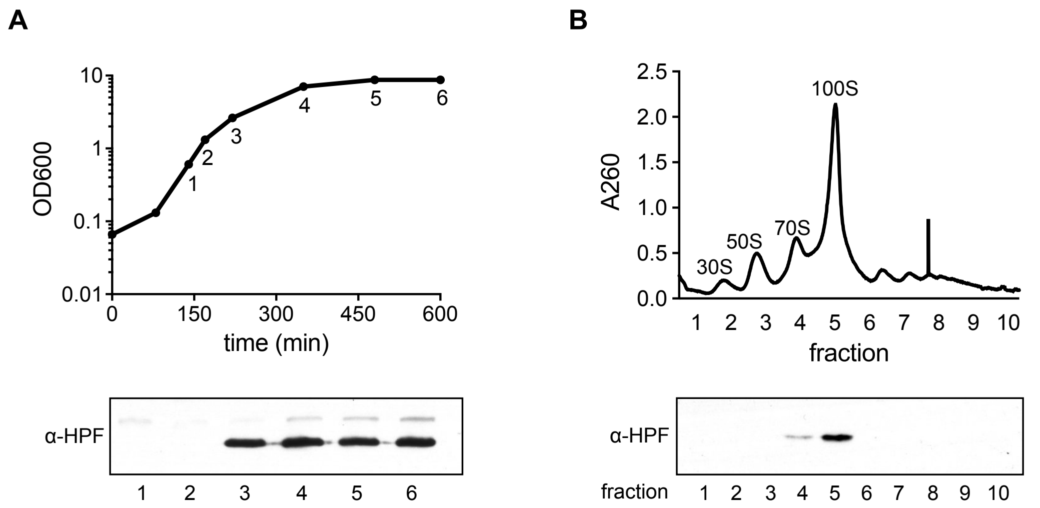


**Fig. S1. HPF is expressed during the transition from log phase to stationary phase.** (A) Growth curve of wild-type *B. subtilis* (JDB1772) with samples harvested at indicated time points for western blot. Polyclonal antibody raised against HPF was used to probe cell lysates. (B) Sucrose gradient sedimentation profiles (10% - 60% sucrose) of ribosomes harvested from sample 3 (~4 hours after inoculation). Gradient fractions were probed with anti-HPF antibody.

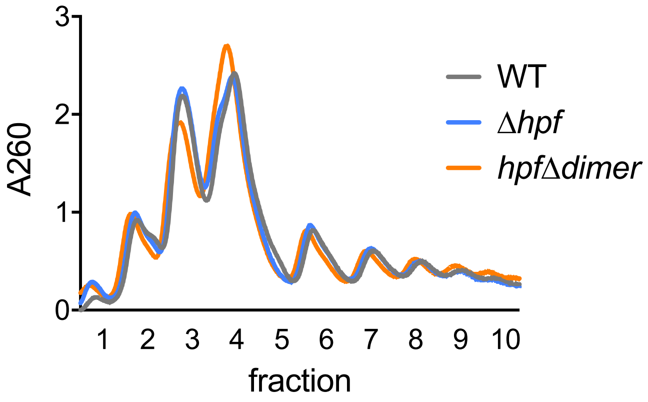


**Fig. S2. Sucrose density gradients of wild-type and HPF deficient strains during log phase.** Wild-type (JDB1772), ∆hpf (JDB4221) and *hpf∆dimer* (JDB4227) strains were grown to late log phase (OD 0.9) and ribosomes were purified from these strains. Ribosome concentrations were normalized by A260 and loaded on a 10%-40% sucrose gradient.

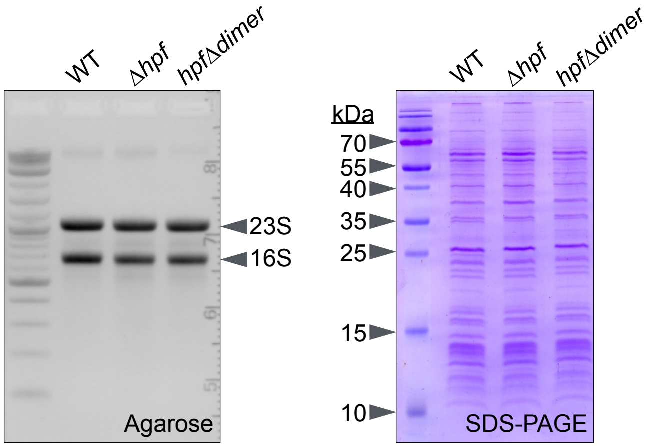


**Fig. S3. Levels of ribosomal RNA and proteins associated with ribosomes from wild-type (JDB1772), Δ*hpf* (JDB4221) and *hpf*Δ*dimer* (JDB4227) strains.** Ribosomal RNA was examined on a 1.5% agarose gel. Ribosomal proteins were resolved on a 15% SDS-PAGE gel and stained with coomassie brilliant blue dye. Asterisks indicate bands the size of wild-type HPF and tagged HPF (HPFΔdimer) protein.

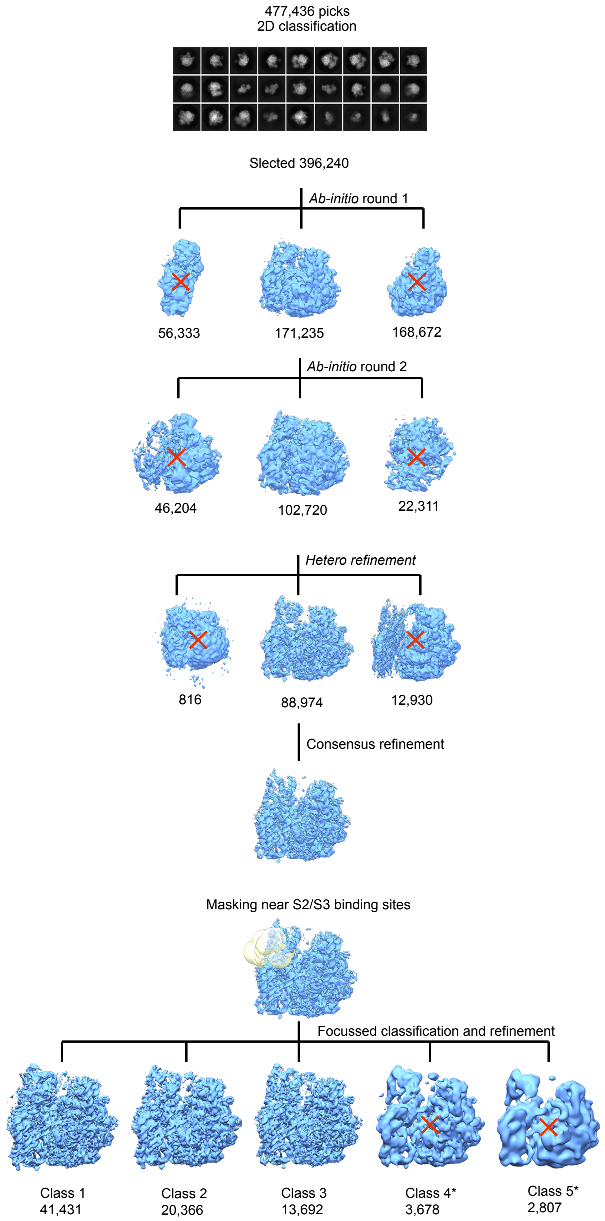


**Fig. S4. Classification and refinement of Δ*hpf* 70S complexes.** After the initial 2D-classification in CryoSPARC, particles were classified in 3D into 30S, 50S, and 70S ribosomal species. 70S ribosomes were selected for another round of *ab-initio* classification, where only ribosomes with well-defined 30S subunit were selected. After the final clean-up using heterogeneous refinement classification, the resulting particle stack was re-refined in RELION to produce "consensus" refinement map. This map was used for focused 3D classification in RELION using the mask generated in the area of interest around the S2 and S3 binding site. Resulting classes were refined in CryoSPARC and analyzed in Chimera. Classes 4 and 5 were poorly resolved and were excluded from further analysis.

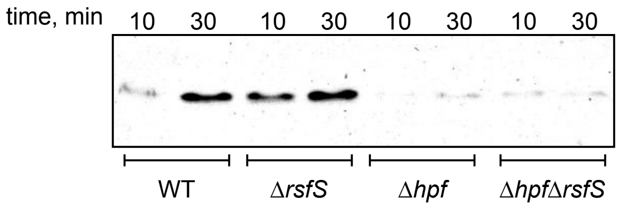


**Fig. S5. *In vitro* translation with strains lacking ribosome silencing factor RsfS.** Ribosomes were purified from wild-type (JDB1772), Δ*rsfS* (JDB4292), Δ*hpf* (JDB4221), or Δ*hpf*Δ*rsfS* (JDB4293) cells grown to stationary phase (~OD 7.0) and added to a ribosome-free *in vitro* translation mix. An M2-tagged template was supplied and reactions were incubated at 37˚C for the indicated times, resolved by SDS-PAGE, and probed with an antibody against M2.

**Table S1. Bacterial Strains and Plasmids**

| Strain | Relevant genotype | Description | Origin |
|---|---|---|---|
| JDB1772 | *trpC2* | B. *subtilis* wild-type parent | Chet Price |
| JDB4221 | *trpC2* Δ*hpf::tet* | *hpf* deletion in wild-type | This study |
| JDB4227 | *trpC2* hpf::*hpf-tag spec* | Flag tagged HPF at endogenous locus | This study |
| JDE3069 | BL21(DE3) pET28b-hpf | For purification of His-HPF This study | |
| JDB4292 | trpC2 Δ*rsfS::kan* | *rsfS* deletion in wild-type | This study |
| JDB4293 | trpC2 Δ*rsfS::kan* ˙Δ*hpf::tet* | *rsfS* deletion in JDB4221 | This study |
| JDE3024 | DH5a pSFP234 | *cotE*-M2 template for in vitro translation | (1) |

**Table S2. Primers used**

| Oligo | Sequence* | Restriction sites | Origin |
|---|---|---|---|
| HF40 | ATATA*GAATTC*AAGGGAGGCGTTCTTT GATGAAC | EcoRI | This study |
| HF41 | ATAT*GCATGC*CGGCCG*GCTAGC*TTCAGTCGGTTCAATTAAGCCATATTTC | SphI and NheI | This study |
| HF38 | TATA*CCATGG*GCATGAACTATAACATCAGAGGAGAAAATATTGAAG | NcoI | This study |
| HF39 | TGGTG*CTCGAG*TTCAGTCGGTTCAATTAAGCCATATTTCC | XhoI | This study |
| HF152 | ATAT*AAGCTT*AGGGAGGCGTTCTTTGATGAAC | HindIII | This study |
| HF177 | CCGG*GCTAGC*TCATTATTCAGTCGGTTCAATTAAGCCAT | NheI | This study |
| SFP169 | CGTAACATAGAGCTTTCCG | | This study |
| SFP170 | GAACAACCTGCACCATTGCAAGAGGATATGTATCTATTTCTC | | This study |
| SFP181 | TTGATCCTTTTTTTATAACAGGAATTCGAGAAGCCTTCCGTGATGTC | | This study |
| SFP182 | CTTCTCCGCTTTTAACGTGC | | This study |
| SFP1 | TCTTGCAATGGTGCAGGTTGTTC | | This study |
| SFP2 | GAATTCCTGTTATAAAAAAAGGATCAA | | This study |

* Restriction sites are in italics

**Dataset S1 (separate file).** Data from mass spectromic analysis of stationary phase ribosomes from wild-type (wt), Δ*hpf* (hpf), and *hpf*Δ*dimer* (hpf-M2). Strains. Normalized strict pept>2 represents normalized protein intensities for each sample. Columns BI – BM show the log2 value of the difference between each mutant and wild-type. 0806 samples were harvested in parallel. 1231 samples were harvested in parallel. Column BK shows the average difference between wild-type and Δ*hpf* ribosomes from the independent 0806 and 1231 samples.
